# Genomic prediction and estimation of marker interaction effects

**DOI:** 10.1101/038935

**Authors:** Lars Rönnegård, Xia Shen

## Abstract

The standard models for genomic prediction assume additive polygenic marker effects. For epistatic models including marker interaction effects, the number of effects to be fitted becomes large, which require computational tools tailored specifically for such models. Here, we extend the methods implemented in the R package bigRR so that marker interaction effects can be computed. Simulation results based on marker data from *Arabidopsis thaliana* show that the inclusion of interaction effects between markers can give a small but significant improvement in genomic predictions. The methods were implemented in the R package EPISbigRR available in the bigRR project on R-Forge. The package includes an introductory vignette to the functions available in EPISbigRR.

**R package URL**: https://r-forge.r-project.org/R/?group_id=1301

## Introduction

Genomic prediction uses whole-genome marker information to predict unobserved phenotypes and is extensively used in livestock breeding. The standard GBLUP model [1] assumes additive polygenic marker effects. The model can be presented either as 1) a linear mixed model with individuals as random effect where the correlation matrix between the random effects is given by the genomic relationship matrix, or as 2) a linear mixed model with independent random marker effects. The generalized ridge regression approach [2] implemented in the bigRR package on CRAN [3] adds additional shrinkage to the marker effects and the shrinkage can vary between markers. Nevertheless, the bigRR package, as well as GBLUP, assumes that the effects are additive.

Interactions between markers may have an important effect on phenotypes and has therefore been suggested to be an important component to be included in models for genomic predictions [4]. Several attempts have been made to perform genomic predictions including epistasis but the direct connection to the fitted marker effects have been largely ignored, most probably because the number of interaction effects becomes very large even for a moderate number of markers.

In this paper, we present a computational tool that computes individual predictions (breeding values) as well as marker interaction effects. The EPISbigRR package is available in the bigRR project on R-Forge. It extends the previously published bigRR package [3] in R.

## Material and Methods

Here, we describe the standard SNP-BLUP and GBLUP models, and how the generalized ridge regression method in the bigRR package [2, 3] can be presented in terms of the SNP-BLUP and GBLUP models. Thereafter, marker interaction effects are introduced.

### SNP-BLUP model

The SNP-BLUP model assumes that the trait *y* (length *n*) is affected by a linear combination of random marker effects *u*. The lengh of *u* is equal to the number of markers, *p*, and the random marker effects are assumed identically and independently distributed (iid) and normal, so that 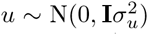. Furthermore, there may be fixed effects ß and the residuals are iid, 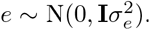. Thus, the linear mixed model is

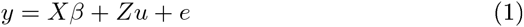

where *X* is a design matrix and *Z* is a scaled incidence matrix for the SNP coding such 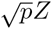 has column means equal to 0 and column variances equal to 1. The estimates of *β* and *u* are computed from the mixed model equations (MME)

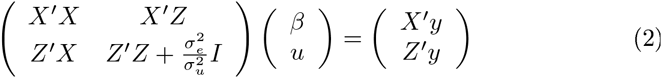

where the variance components, 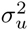 and 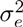 are estimated using REML. Thus, from this model we can compute the fitted effect of the *i*:th marker, *û_i_*, and its hat value *h_ii_* [2].

### GBLUP model

The GBLUP model is equivalent to the SNP-BLUP model. Here, the individual random effects *a* (length *n*) are defined such that *a* = *Zu*, and we thereby have:

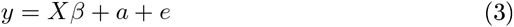

where 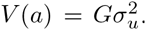. Here *G* is the genomic relationship matrix and we have *G* = *ZZ′*. Thus, the MME are

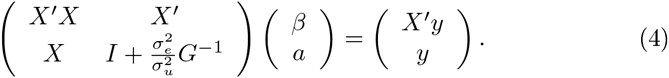

Shen et al. [2] showed how the *û* can be computed given the fitted values *â*, and also how the hat values can be transformed between the two models.

### Generalized ridge regression implemented in the bigRR package

The generalized ridge regression model implemented in the bigRR package allows variable shrinkage for different markers by introducing a diagonal matrix Λ such that 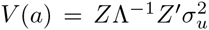 in the GBLUP model, and the mixed model equations for the SNP BLUP model is

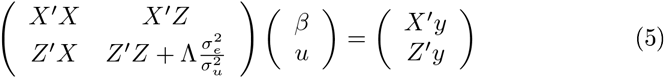

The generalized ridge regression model in bigRR computes the diagonal elements of Λ as

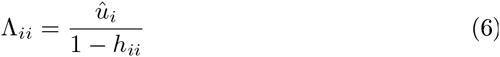

and it is easy to compute the fitted marker effects *û* from the fitted individual effects *â* making the computations fast.

### Epistasis

Below we present the SNP-BLUP interaction model corresponding to a GBLUP model that uses a direct Hadamard product to compute the correlation between individual random effects. We show that this model actually includes dominance effects, and explain how the correlations due to dominance and marker interaction effects can be separated.

### SNP-BLUP model including marker interaction effects

The SNP-BLUP model can be extended to include marker interaction effects [5]

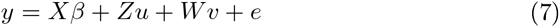

*v* marker interaction effects, and the matrix *W* is constructed so that *W_j_* = *Z_i_* ⊙ *Z* with subscript giving column index with *j* = (*i* — 1)*p* + *i* where *p* is the number of columns in *Z* and ⊙ is the direct Hadamard product. Thus, *W* has *n* rows and *p* × *p* columns.

### GBLUP model including epistasis

The equivalent GBLUP model is

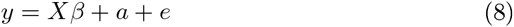

with 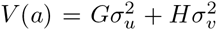. By putting *H* = *G* ⊙ *G*, the GBLUP model including epistasis becomes equivalent to the SNP BLUP model including marker interaction effects.

It should be noted though, that the way we have define *W* it actually also includes dominance effects, and so does the GBLUP model when we let *H* = *G* ⊙ *G*. This is because *W* includes interactions effects between column *i* and *i*, ie a dominance effect.

### Excluding dominance effects from the interaction effects model

In eq. (7), *W* was constructed such that it includes the pair-wise interactions twice and also the dominance interaction effects. As Xu [5] points out, however, the columns in *W* should be constructed so that W_j_ = *Z_i_* ⊙ *Z*_(*i*+1):*p*_. Hence, the interaction of a locus with itself (i.e. dominance) should not be included and each pair-wise interaction is only accounted for once. The equivalent covariance matrix for the epistatic effects is constructed as

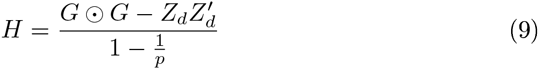

where *Z_d_* = *Z* ⊙ *Z* is the model matrix for the dominance effects. As the number of SNPs is typically very large, i.e. large *p*, the denominator can be ignored.

### What has been added to EPISbigRR?

In EPISbigRR the extra shrinkage is computed from the estimates of *v* and their associated hat values, such that 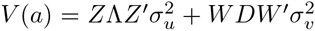 where D is a diagonal matrix with diagonal elements 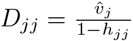 for interaction effect *j*.

In order to fit the epistatic effects model (7), EPISbigRR includes some new features compared to bigRR: more than one variance component can be fitted, and the shrinkage can be controlled separately for the different variance components. The hugeRR and hugeRR^update functions stores matrices temporarily using the DatABEL format of the GenABEL [6] which makes the computations feasible for a large number of markers. The EPISbigRR package includes a vignette illustrating the available functions.

## Results

The Arabidopsis data including 84 individuals available in the bigRR package was used to evaluate the advantage of including epistasis to genomic predictions. Estimates of marker interaction effects are found in the EPISbigRR package vignette, and here we focus on the prediction accuracy of the genomic predictions.

500 cross-validation replicates where performed where 70 out of the 84 phenotypes were sampled for the training set and 14 for the test set. Phenotypes were simulated for the parameters 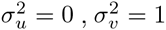 and 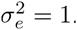. The epistatic GBLUP model was compared to the ordinary GBLUP model, and the comparsion was simply made using the hglm function [7]. For 276 out of the 500 replicates the epistatic GBLUP model outperformed the ordinary GBLUP model. A small but significant improvement (P = 0.009).

## Discussion

We have developed a tool that makes it computationally feasible to compute marker interaction effects for genomic predictions. An interesting future development, which seems rather straightforward, would be to parallelize the computations. This should be rather easy since the computations are performed in parts and intermediate results stored in seprate files (in DatABEL format) using the hugeRR and hugeRR_update functions.

We have also clarified the difference between the model specified by Xu [5] and the epistatic GBLUP model using *G* ⊙ *G* as covariance matrix for the epistatic effects (e.g. [4]). The correct covariance matrix to be used is easily constructed as *G* ⊙ *G* – 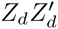 where *Z_d_* is the scaled model matrix for the dominance effects (see eq. (9)).

For the interaction effects model, the number of pair-wise marker effects to be fitted becomes enormous. Our tool makes these computations feasible in time but since we have an extreme n≪p problem the effects are fitted with great uncertainty and the generalized ridge regression estimates will be sensitive to the hat values being close to 1.

Hill [8] and Crow [9] explain that epistasis will have no substantial effect on the response from recurrent selection, because the genetic gain induced by epistasis arises from the gametic disequilibrium among the epistatic loci. Nevertheless, the importance of epistasis is still a matter of debate [10] and is expected to be so far into the future.

The developed package should be useful for further studies of the importance of epistasis in genomic predictions.

